# Gene-by-environment interactions influence the fitness cost of gene copy-number variation in yeast

**DOI:** 10.1101/2023.05.11.540375

**Authors:** DeElegant Robinson, Elena Vanacloig-Pedros, Ruoyi Cai, Michael Place, James Hose, Audrey P Gasch

**Author notes:** Corresponding author: Audrey P. Gasch. Ann & Robert H. Lurie Children’s Hospital of Chicago, Chicago IL 60611. Department of Biostatistics, University of Washington, Seattle, WA, 98195. Both authors contributed equally to this work (DeElegant Robinson, Elena Vanacloig-Pedros).

## Abstract

Variation in gene copy number can alter gene expression and influence downstream phenotypes; thus copy-number variation (CNV) provides a route for rapid evolution if the benefits outweigh the cost. We recently showed that genetic background significantly influences how yeast cells respond to gene over-expression (OE), revealing that the fitness costs of CNV can vary substantially with genetic background in a common-garden environment. But the interplay between CNV tolerance and environment remains unexplored on a genomic scale. Here we measured the tolerance to gene OE in four genetically distinct *Saccharomyces cerevisiae* strains grown under sodium chloride (NaCl) stress. OE genes that are commonly deleterious during NaCl stress recapitulated those commonly deleterious under standard conditions. However, NaCl stress uncovered novel differences in strain responses to gene OE. West African strain NCYC3290 and North American oak isolate YPS128 are more sensitive to NaCl stress than vineyard BC187 and laboratory strain BY4743. Consistently, NCYC3290 and YPS128 showed the greatest sensitivities to gene OE. Although most genes were deleterious, hundreds were beneficial when overexpressed – remarkably, most of these effects were strain specific. Few beneficial genes were shared between the NaCl-sensitive isolates, implicating mechanistic differences behind their NaCl sensitivity. Transcriptomic analysis suggested underlying vulnerabilities and tolerances across strains, and pointed to natural CNV of a sodium export pump that likely contributes to strain-specific responses to OE of other genes. Our results reveal extensive strain-by-environment interaction in the response to gene CNV, raising important implications for the accessibility of CNV-dependent evolutionary routes under times of stress.

## INTRODUCTION

Many unicellular organisms like budding yeast *Saccharomyces cerevisiae* live in environments that fluctuate. Yeast cells can exist in a range of habitats, from fruits and trees to insects and human-associated niches (1, 2). Many of these habitats vary over time and space. Sudden environmental changes occur frequently in nature and can include fluctuations in nutrient availability, temperature, exposure to toxins, and other conditions (3). Thus, cells have evolved to deal with changing environments including changes that are stressful. Genetic variation has a substantial influence on stress tolerance and response, due in part to neutral genetic drift but also influenced by adaptive changes. For example, strong selective pressure for copper exposure, rapid freeze-thaw cycles, dessication tolerance, and other conditions are thought to have led to selection for specific genetic backgrounds (4-7). A major focus in evolutionary biology has been to understand modes of evolution and the genetic architecture of differences in environmental tolerance. While single-nucleotide changes can influence phenotype, differences in gene copy number provide a driving force, especially in stressful environments (8-12). CNV can impart an immediate effect on gene expression, which in turn can have an immediate influence on phenotype; if the benefits outweigh the costs, CNVs can become fixed (8, 13-17). Yet how the cost and benefit of CNV varies with genetic background is only beginning to emerge.

We previously showed that the fitness consequences of gene overexpression (OE), used to model CNV, can vary substantially depending on genetic background. We identified shared and unique responses to each of ∼4,700 yeast genes expressed on a high-copy plasmid in 15 different strains of *S. cerevisiae* selected from diverse niches and locations from around the globe (18). This library expresses each gene from its native regulatory sequences, along with a unique DNA barcode that can be quantified by sequencing; relative fitness of each gene can be inferred by changes in barcode abundance after competitive growth compared to the starting library. While amplification of >400 genes is commonly deleterious to many strains, the majority of fitness effects were seen in only a subset of strains. This reveals that the fitness effects of gene OE vary widely with host genome, implicating extensive strain-by-CNV interactions. This implies that that different genetic backgrounds will have differential access to evolutionary routes that involve CNV; indeed, strains exposed to extreme selection evolve through different mechanisms, including those that leverage CNV and those that do not (19-23).

A major remaining question is how the environment influences genetic variation in the response to CNV. In nature, cells can experience many different conditions and environments; thus, understanding genotype-environment interactions (GxE) on the consequences of CNV is important (24-26). Here we explored this GxE relationship by examining how the response to gene OE varies across strains grown in a stressful condition. We chose sodium chloride (NaCl) as a stress because of the wealth of molecular information on how yeast cells respond to NaCl stress and how cells regulate the response (27, 28). Exposing yeast cells to NaCl causes osmotic stress and ion toxicity, which provoke diverse downstream effects including rapid water efflux, increased Na^+^ influx and concentration in the cytosol, production of internal osmolytes among other metabolic changes, and mobilization of transcriptomic changes, including activation of the environmental stress response (ESR) (29-33). Several signaling pathways are known to respond to NaCl stress, including High Osmolarity Glycerol (HOG) pathway, AMP-responsive kinase Snf1, and Calcineurin (28, 34-38). These pathways can mediate defense strategies such as regulating glycerol accumulation, protecting against protein misfolding, and mediating metabolic changes.

To explore how GxE interactions influence the consequences of gene CNV, we expressed the MoBY 2.0 gene OE library (39, 40) in four different yeast strains, each grown for 10 generations in 0.7M NaCl. While many of the detrimental effects were shared across strains, most of the beneficial OE genes were strain-specific. Transcriptomic and genomic analysis revealed several important features of strain-specific responses to NaCl and gene CNV, which may translate to strain-specific evolutionary trajectories during times of stress.

## METHODS

### Strains and growth conditions

Strains used in this study include the laboratory strain BY4743 (MATa/α his3*Δ*1/his3*Δ*1 leu2*Δ*0/leu2*Δ*0 LYS2/lys2*Δ*0 met15*Δ*0/MET15 ura3*Δ*0/ura3*Δ*0), BC187 (41), NCYC3290 (42), and YPS128 (43). Strains were grown in rich YPD medium (10 g/L yeast extract, 20 g/L peptone, 20 g/L dextrose) with G418 (200mg/L) for plasmid selection in shake flasks at 30°C, with or without 0.7M NaCl. As described previously (18), strains were transformed with the high-copy MoBY 2.0 library (39, 40) to at least 5-fold replication (∼25,000 transformants per strain and library of ∼5,000 unique plasmids). Colonies isolated on plates from the transformation were scraped, pooled, and stored at -80°C.

### Fitness measurements

The competition experiments were performed as previously described (40). Frozen library-transformed stocks were thawed and placed in 100ml of liquid YPD with 0.7M NaCl and G418 (200mg/L), at a starting OD_600_ of 0.05. Cultures were transferred to fresh media with or without appropriate supplementation after 5 generations to keep cells in log phase. Cells were harvested after 10 generations and cell pellets were stored at -80°C.

### Barcode sequencing and analysis

Plasmids were collected from each culture aliquot using QIAprep spin miniprep kits (Qiagen, Hilden, Germany). The pool of barcodes was amplified as previously described (2). Samples were pooled, split, and sequenced across three lanes of an Illumina HiSeq Rapid Run using single end 100bp reads. Sequencing data are available in the NIH GEO database under accession number GSE226247-8.

The data were normalized using library-size normalization as described in (18), with one modification: some barcodes repeatedly rose to very high read count in the wild strains. To avoid these genes skewing the library normalization, we calculated total sample read counts excluding genes with >50,000 reads, then divided all read counts (including these highly abundant counts) by that normalization factor. Some highly deleterious genes completely drop out of the population after NaCl outgrowth; for these genes we imputed missing data similarly to what was described previously (18) as follows: for genes that were well measured at the starting point (>20 normalized read counts in all three replicates) but missing after NaCl outgrowth, we added one pseudocount at generation 10. After library normalization, we scaled all values by 1,000,000 and rounded to the nearest integer for *edgeR* analysis (44). The processed and normalized data (available in Dataset 1) were used as input to *edgeR* using a linear model with generation and strain as factors. Genes with FDR < 0.05 were considered significant (45); the output from edgeR is provided in Dataset 2. Relative fitness scores were calculated as the log_2_ ratio of normalized read counts after versus before outgrowth. Hierarchical clustering (46) was performed using Cluster 3.0 and visualized using Java TreeView (47). Data for YPD rich media without NaCl were taken from (18).

We considered genes with a fitness benefit as those with a significant positive fitness effect (FDR < 0.05); because many genes selected in BY4743 and BC187 strains had very small effect sizes, we also applied a magnitude threshold, requiring a log_2_ fitness effect of at least 0.8564, which was the smallest beneficial fitness effect of significant genes (FDR < 0.05) in NCYC3290 and YPS128. 314 genes met these criteria in at least one of the four strains analyzed (Dataset 3). Functional and biophysical enrichments were evaluated using Hypergeometric tests, taking p-value ≤ 10^−4^ as significant.

### Transcriptome profiling and analysis

Yeast strains were grown in biological triplicate in rich YPD medium at 30°C with shaking, for three generations to an OD600 ∼0.5; all strains were grown in parallel for each replicate, allowing paired downstream analysis. Cells grown in rich medium were shifted to media with 0.7M NaCl, and samples were collected before and at 30 minutes and 3 hours after the shift. Cells were collected by centrifugation, flash frozen, and maintained at -80°C until RNA extraction. Total RNA was extracted by hot phenol lysis (48), digested with Turbo DNase (Invitrogen) for 30 min at 37°C, and precipitated with 5 M lithium acetate for 30 min at -20°C. rRNA depletion was performed using the Ribo-Zero (Yeast) rRNA Removal Kit (Illumina, San Diego, CA) and libraries were generated according to the TruSeq Stranded Total RNA kit and purified using a Axygen AxyPrep MAG PCR Clean-Up Kit (Axygen). The samples were pooled, re-split, and run across three lanes on an Illumina HiSeq 2500 sequencer, generating single-end 100 bp reads, with ∼7,494,848 reads per sample. Sequencing data are available in the NIH GEO database under accession number GSE226246.

Reads were processed using Trimmomatic version 0.3 (49), and mapped to the S288c reference genome (version R64-1-1) with bwa-mem (version 0.7.12-r1039) (50). Read counts for each gene were calculated by HT-Seq (version 0.6.0) (51) and normalized using the TMM method in *edgeR* (44). We used a linear model in *edgeR* with strain background as a factor and paired replicates, identifying genes differentially expressed in each strain relative to the average of all strains taking FDR <0.05 as significant. Hierarchical clustering (46) was performed by Cluster 3.0 and visualized using Java TreeView (47). Functional analysis and enriched GO categories for each sample were obtained using hypergeometric tests, taking p-value ≤ 10^−4^ as significant. *EdgeR* identified 1,114 genes whose fold-change in expression after NaCl was different in at least one strain compared to the mean fold-change across strains (FDR < 0.05, Dataset 4, Tab 1). Relative expression was represented as the log_2_ (fold change) in each strain, comparing normalized read counts 30 min or 3h versus before NaCl treatment in each strain. Comparing basal (unstressed) expression across strains identified 608 genes whose expression was significantly different from the mean (FDR<0.05) in at least one of the strains (Dataset 4, Tab 2).

### ENA gene CNV analysis

We analyzed previously published DNA-seq data generated for these strains and mapped to the S288c reference sequence (42, 52, 53). We normalized read counts at all positions to the genomic media read count, and then randomly selected three representative genes from three different chromosomes (YDR229W, YGR125W, YJL059W) whose DNA-seq read abundance was at the median read depth of the genome. These genes were taken to represent genes at single copy per haploid genome. To avoid mapping errors to the S288c reference that contains multiple highly-similar ENA genes, we re-mapped reads to a reference sequence consisting of S288c *ENA1* (*YDR040C*) and the three representative single-copy genes using bwa-mem (50) and then used samtools mpileup function (54) to plot the coverage of each base pair in the reference sequence. Read counts were normalized to the median coverage of the three reference genes. Fig 5 shows the running average of normalized read count over 500bp windows.

## RESULTS

To investigate the effects of gene OE under NaCl stress, we focused on a subset of diploid strains analyzed in a previous study from our lab, selecting three strains representing distinct genetic lineages: West African strain NCYC3290, vineyard strain BC187, and North American oak isolate YPS128, along with common lab strain BY4743 as a well-studied reference (42, 55). In addition to genetic differences, these strains display extensive phenotypic diversity in several different environmental conditions, including osmotic stress (53, 55-57). We first measured strain growth rates in 0.3M and 0.7M NaCl to characterize NaCl tolerance. Growth of all strains was reduced in NaCl compared to rich medium (Fig 1). However, there was significant variation in strain responses. West African NCYC3290 and to a lesser extent oak soil YPS128 strains were much more sensitive to the higher dose of NaCl, indicated by their 8.5X and 3.5X reduction in doubling time compared to only ∼2X reduction in vineyard strain BC187 and lab strain BY4743. We chose the higher dose of NaCl to interrogate how this environment influences strain-specific responses to CNV.

**Figure 1:**
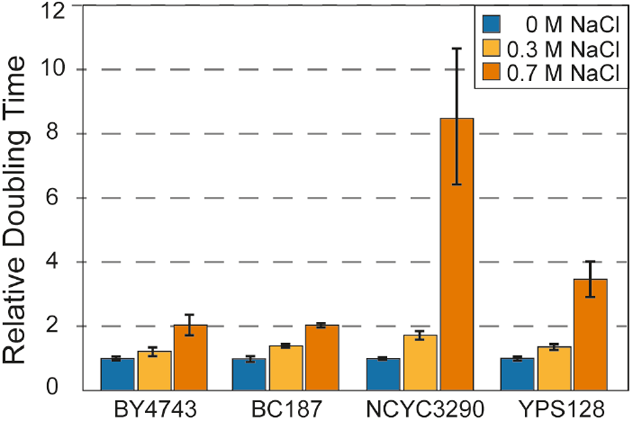
Strains vary in salt sensitivity. Average and standard deviation (n=3) of doubling times of each strain grown in the absence or presence of NaCl according to the key, normalized to the unstressed growth rate of that strain.

Next, we quantified the fitness effects of gene OE across different genetic backgrounds when cells were subjected to a stressful environment of 0.7M NaCl. Each strain was transformed with the library, which includes ∼5,000 genes cloned from S288c along with their native upstream and downstream sequences, cloned onto a 2-micron replication plasmid (39, 40). Each plasmid also carries a DNA barcode, which can be identified and quantified through deep sequencing of the pooled library. We note that one limitation of this approach is that some genes that are not expressed in salt conditions will not show increased expression and thus will be scored as neutral. Cells were inoculated into rich medium and an aliquot collected before (‘0 generation’ sample) and after 10 generations of growth in 0.7M NaCl, in biological triplicate. Gene abundances before and after outgrowth were identified by deep sequencing of plasmid barcodes.

Past analysis showed that these three wild strains carry 2-3 copies of the 2-micron plasmid from this library, per haploid genome, making results from these strains directly comparable. The laboratory strain is distinct from many wild strains and carries ∼11 copies per haploid genome (18). To account for these differences, we normalized sequencing data for each library and calculated the log_2_ ratio of normalized barcode read counts after NaCl treatment *versus* before (see Methods). This reflects the relative fitness cost of each gene compared to the library expressed in that strain. Plasmids that carry genes that are detrimental when OE drop in frequency in the population, either because of reduced cell growth in the population or because cells suppress the abundance of toxic plasmids (58), both of which we interpret as a relative fitness defect. In contrast, beneficial plasmids will rise in frequency in the population over time. We used linear modeling to identify genes with a significant fitness effect in each strain (FDR < 0.05).

### Common and unique fitness consequences across strains and environments

We identified a total of 3,644 genes (from 4,133 interrogated) whose OE produced a relative fitness effect in at least one strain growing under NaCl stress (FDR< 0.05, Fig 2). Yet the number and magnitude of fitness costs varied substantially. Lab strain BY4743 showed the least impact of gene OE, both in terms of number of deleterious OE genes and their impact on fitness (Fig 2A-B), followed by BC187 which also showed relatively mild fitness effects for most genes. For both of these strains, the distribution of relative fitness costs was similar with and without NaCl, consistent with their ability to grow well in salt-containing medium (Fig 1). In contrast, the NaCl-sensitive strains showed several key differences. First, both strains were more sensitive to gene OE in the absence of stress, indicated both by the number of deleterious OE genes and their greater effect sizes compared to NaCl-tolerant strains. Second, both strains – but especially the most NaCl-sensitive, NCYC3290 – were much more sensitive to gene OE in the presence of NaCl. For example, the vast majority of OE genes were deleterious in NCYC3290 and with severe fitness costs (Fig 2A-C).

**Figure 2:**
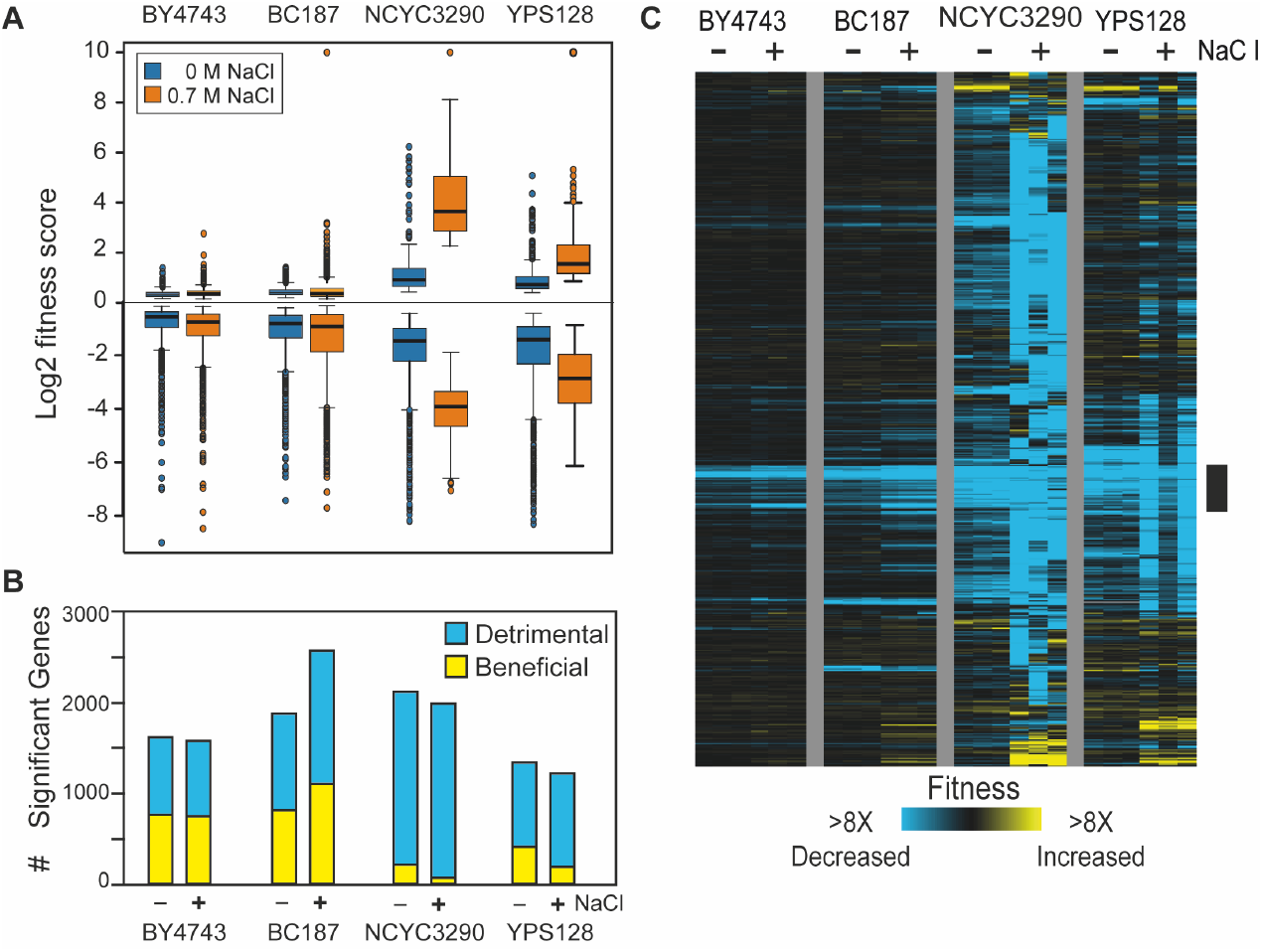
Fitness consequences vary by strain. A) Distribution of average log_2_ fitness scores in each strain grown 10 generations in rich medium in the absence of NaCl (blue) or in medium with 0.7M NaCl (orange). B) The number of detrimental (blue) or beneficial (yellow) genes in each strain and media condition. C) Hierarchical clustering of 3,644 genes with a fitness effect (FDR < 0.05) in at least one strain grown in NaCl stress. Each row is a specific gene, each column is a biological replicate of that strain grown in the absence (-) or presence (+) of NaCl. Blue and yellow values represent genes that dropped or rose in abundance during competitive growth, reflecting decreased or increased fitness effects according to the key. The black bar indicates a cluster of genes whose OE is detrimental to varying degrees in all strains and under both conditions. Data from NaCl-free conditions were taken from (18).

To explore the patterns of fitness costs across strains and conditions, we hierarchically clustered OE genes that had a relative fitness effect in at least one strain. The resulting heat map illustrates the commonalities and differences across strains and conditions. We identified one cluster of ∼200 OE genes (Fig 2C, black bar) that were deleterious in all four 4 strains in both YPD and NaCl stress conditions, to varying degrees. Consistent with past work from our lab (18), this gene group was heavily enriched for genes involved in translation, including ribosomes and ribosome biogenesis factors, protein folding factors, and genes repressed during stress in the Environmental Stress Response (p < 10^−4^, Hypergeometric tests). But even for these commonly deleterious OE genes the magnitude of the effect varied, especially for the wild strains where the fitness cost was more severe in the presence of NaCl. These results are consistent with the notion that the cost of gene OE is greater in strains already experiencing suboptimal conditions (see Discussion).

### Beneficial genes vary widely across strains

We were particularly interested in genes whose OE provides a benefit under NaCl stress, since these may provide adaptive value. We identified and hierarchically clustered 331 genes whose OE produced a significant benefit in NaCl over an effect threshold in at least one strain (Fig 3A, see Methods). BY4743 and BC187 had a substantial number of genes that passed our statistical threshold but were of very small effect size (hence the use of a magnitude threshold, identifying only 23 beneficial OE genes for BY4743). In contrast, 116, 65 and 183 genes met our criteria of providing a benefit to BC187, NCYC3290 and YPS128, respectively. Interestingly, there was only small overlap in which genes were beneficial during NaCl stress, even for the two NaCl-sensitive strains (Fig 3B). This strongly implies that the genetic basis for NaCl sensitivity is different in the two sensitive strains.

**Figure 3.**
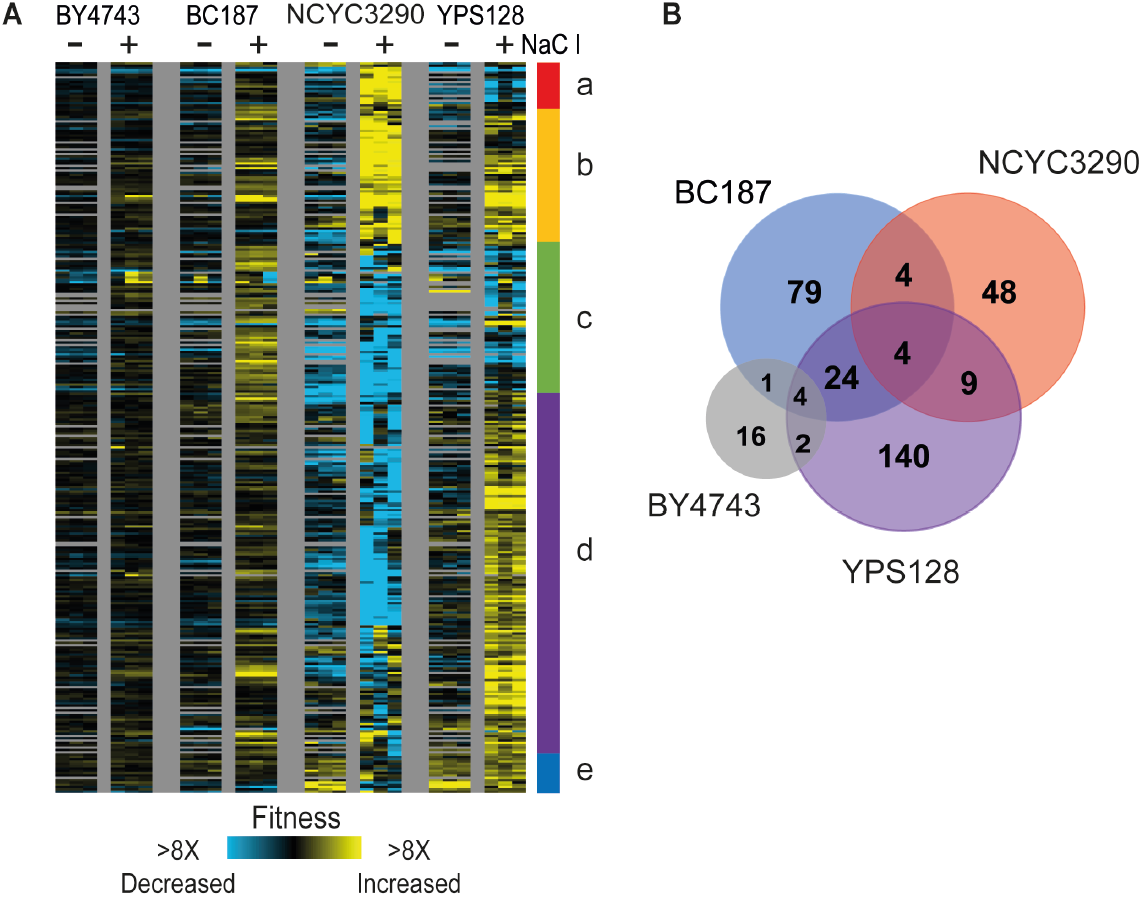
Beneficial genes vary widely by strain. A) Hierarchical clustering of 314 genes that were beneficial in at least one strain (see Methods), as described in Figure 2; grey values indicate missing values. Several clusters were enriched for functional groups (p<1×10^−4^, Hypergeometric test) including genes involved in a) cell-cycle entry, c) negative regulation of heterochromatin, d) cytoplasmic mRNA processing body assembly, snoRNA processing. B) Overlap of beneficial genes identified in each strain.

Only four genes were scored as beneficial to all three wild strains but not BY4743 (Fig 3B). These included calmodulin kinase *CMK2*, Snf1-related kinase *HAL5* that regulates ion tolerance (59, 60), CK2 kinase subunit *CKA1* that has been implicated in NaCl stress (61, 62), and *GCD10* that encodes a tRNA methyl transferase. It is interesting that three of the four genes are kinases that could have a wide range of downstream effects on physiology. Expanding to genes shared between the two sensitive strains identified several other genes involved in cation homeostasis including sodium and other-cation transporter *QDR2*, kinase *SAT4* involved in sodium tolerance (60), and diacylglycerol kinase *DGK1* (itself a target of CK2 (63)). This group was also enriched for genes regulated by the HOG-regulated osmotic stress transcription factor Sko1 (p=5×10^−4^, Hypergeometric test), which is interesting in the context of the NaCl sensitivity of these strains. Interestingly, the total set of genes whose OE was beneficial to YPS128 were enriched for genes involved in mRNA P-body and stress granule assembly and ergosterol biosynthesis (p<1×10^−4^, Hypergeometric test). BC187 also benefited from OE of many RNA binding proteins, several of which were shared with YPS128. In contrast, genes beneficial to NCYC3290 were enriched for those involved in glycosyl-group transferase activity and flocculation. Together, these results indicate that the fitness consequences of beneficial genes are largely strain specific, in some cases causing opposing fitness effects in different strains (Fig 3, clusters a, c, and d).

### Transcriptomic analysis implicates genetic modifiers

The low overlap in beneficial OE genes among the NaCl sensitive strains suggests that NaCl sensitivity is explained by different genetic or physiological limitations. In attempt to better understand background-specific effects that could influence these differences, we characterized each strain’s transcriptomic response to NaCl shock. Cells grown in rich medium were shifted to media with 0.7M NaCl, and samples were collected before and at 30 minutes and 3 hours after the shift, in biological triplicate (see Methods). We chose these timepoints to explore the response to NaCl immediately after acute shock and at a later timepoint that represents the acclimated state. We expected that the sensitive strains may show substantial differences in transcriptomic response to the shock, reflecting their increased NaCl sensitivity; however this was not the case. Using a linear model to identify strain-by-environment interactions, we identified 1,141 genes whose expression to NaCl differed across strains at one or both time points after shock (FDR < 0.05, see Methods). The four strains showed fairly similar gene expression changes at both induced and repressed genes (Fig 4A) – there were no large gene groups that were uniquely induced or repressed in any of the strains. Furthermore, the magnitude of many of those gene expression changes were similar across strains. This was especially surprising for the sensitive strains, since we expected that they may exhibit larger magnitude changes if they experience a stronger stress from 0.7M NaCl treatment.

**Figure 4.**
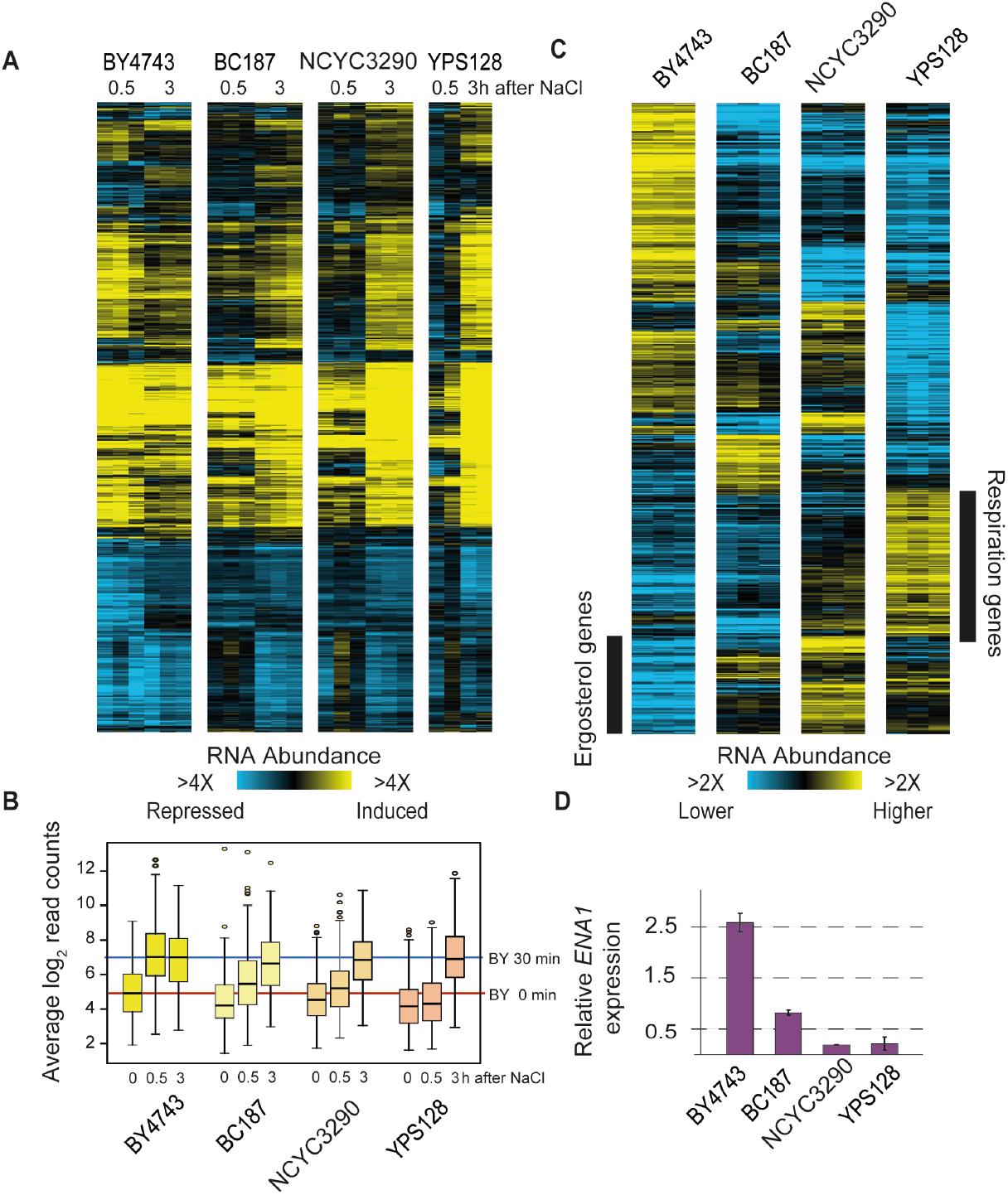
Gene expression differences implicate strain-specific physiology. A) Hierarchical clustering of 1,141 genes whose log_2_(fold change) in response to NaCl was different from the mean in at least one of the four strains (FDR < 0.05). As shown for Figure 2, except that here values represent relative mRNA abundance in stressed samples versus the prestressed sample from that strain. Each column represents one of three biological replicates (or two in the case of YPS128). B) The distribution of log_2_ normalized read counts for genes induced in the ESR across strains and time points. The median abundance of genes in the laboratory strain before and at 30 min after NaCl treatment is indicated with red and blue lines. C) Hierarchical clustering of 608 genes whose basal expression before NaCl is different from the mean in at least one of the four strains (FDR < 0.05). Each column is one of three biological replicates of cells growing in rich medium in the absence of stress, where log_2_ values represent expression relative to the mean of all strains according to the key. D) Average and standard deviation of *ENA1* expression in the absence of stress relative to the mean of all strains.

Despite these global similarities, on closer inspection we noticed a difference in the timing of the response, especially compared to the BY4743 laboratory strain. The lab strain had much larger changes in expression at 30 min after NaCl shock than the other strains, and many of these expression differences were already subsiding by 3 hours. Genes induced in the ESR provide a good representation (Fig 4B): these genes show much larger and earlier expression changes in the laboratory strain. In contrast, all three wild strains showed delayed expression changes, of genes induced in the ESR (Fig 4B) and other genes more broadly (Fig 4A). Much of the transcriptome response is regulated by the Hog1 kinase that responds to osmotic stress. Interestingly, it is well known that the timing of Hog1 signaling varies with the dose of ionic stress: at higher doses of stress, signaling is delayed and produces a delayed transcriptomic response (64-67). This suggested that wild strains could all be experiencing a higher level of NaCl stress that delays the transcriptomic response (yet does not grossly change the magnitude or genes in the response, see Discussion). There was little relationship between genes expression and gene fitness benefits in each strain (aside of YPS128, in which seven beneficial genes were more highly induced in that strain upon NaCl treatment). In fact, only 20% of genes beneficial in one or more strain (Fig 3B) were differentially expressed in any strain (Fig 4A), and nearly half of those (43%) were repressed by NaCl across strains. This is not entirely surprising, since most genes with expression changes during NaCl treatment have no bearing on surviving NaCl treatment, and many genes important for NaCl survival are actually repressed at the transcript level (68-70).

Although the transcript changes to NaCl were not wildly different across strains aside of the timing, we wondered if basal expression differences in the strains, before NaCl exposure, could be informative. We therefore identified 608 genes whose expression was significantly different in at least one strain compared to the mean (FDR < 0.05, see Methods). Here, expression differences were more noticeable across the strains (Fig 4C). In fact, the laboratory strain was a clear outlier compared to the three wild strains: one large group of genes was expressed significantly higher in BY4743 in the absence of stress, and this group was enriched for stress-defense genes (see also Fig 4B), oxidoreductases, amino acid biosynthesis genes, transporters, and genes encoding proteins localized to the membrane and vacuole (p<1e-4, Hypergeometric test). This result suggests that BY4743 is already prepared for stress even before exposure (see Discussion). A second group of genes was expressed significantly lower in the lab strain, and these were heavily enriched for genes encoding respiration factors and ergosterol biosynthesis genes and targets of Hap1 (Fig 4C). Some of these differences may result from known polymorphisms in S288c-derived strains that affect the regulation of those genes (including *MKT1, MIP1, HAP1* along with auxotrophic markers that affect mitochondrial and respiratory functions and/or ergosterol-gene expression (71-73)).

Remarkably, among the genes with much higher basal expression in BY4743 were several linked directly to Na+ transport, including *ENA1, ENA5*, and *NHA1*. In fact, *ENA* genes coding for P-type ATPase sodium pumps are known to have undergone tandem duplications in different strains including BY4743, which harbors three copies per haploid genome, each with high sequence homology (74-78). *ENA* genes are important components in the saline detoxification (although they are not present in the MoBY 2.0 library), and strains with higher *ENA* copy number are well known to have correspondingly higher sodium tolerance (56, 79-82). We plotted the relative abundance of RNA-seq reads mapping to *ENA1* as a representative and found that expression varied wildly across strains. The two sensitive strains showed very low expression of *ENA1* relative to the other strains; expression in vineyard strain BC187 was ∼4X higher than NCYC3290, while expression in lab strain BY4743 was 14X higher (Fig 4C). These differences raised the possibility of underlying gene copy-number differences.

To investigate if underlying expression differences were due to natural CNV of these genes, we interrogated DNA-seq reads mapped to a single ENA gene copy relative to three representative genes present in single copy per haploid genome (see Methods). NCYC3290 and YPS128 both showed per-base coverage that was similar to the single copy genes, barring some fluctuation in read depth that may be due to polymorphism-dependent mapping errors: the median read count for *ENA1* versus the median of the single-copy genes was 1.1 and 1.0 for NCYC3290 and YPS128, respectively (Fig 5A). In contrast, BC187 showed read distribution that was ∼3-4X higher across most of the gene, excluding the very amino terminus (median *ENA1* read count versus single-copy genes of 3.1). As a control for our analysis method, we plotted reads for a related lab strain, W303, that was sequenced with the same pipeline as the wild strains – the median read depth of *ENA1* versus single-copy genes was 3.7, consistent with the known four *ENA* copies in this strain (83). The reduced coverage at the ends of the gene could be due to incomplete gene duplication, or it may reflect substantial polymorphisms between BC187 and the reference genome that obscure copy number. The latter possibility is supported by the known high rate of evolution of ENA genes, including at least one case of introgression from *S. paradoxus* (56, 75, 79, 81). We conclude that BC187 has 3 ENA gene copies per haploid genome, although the functionality of all copies remains to be explored.

**Figure 5.**
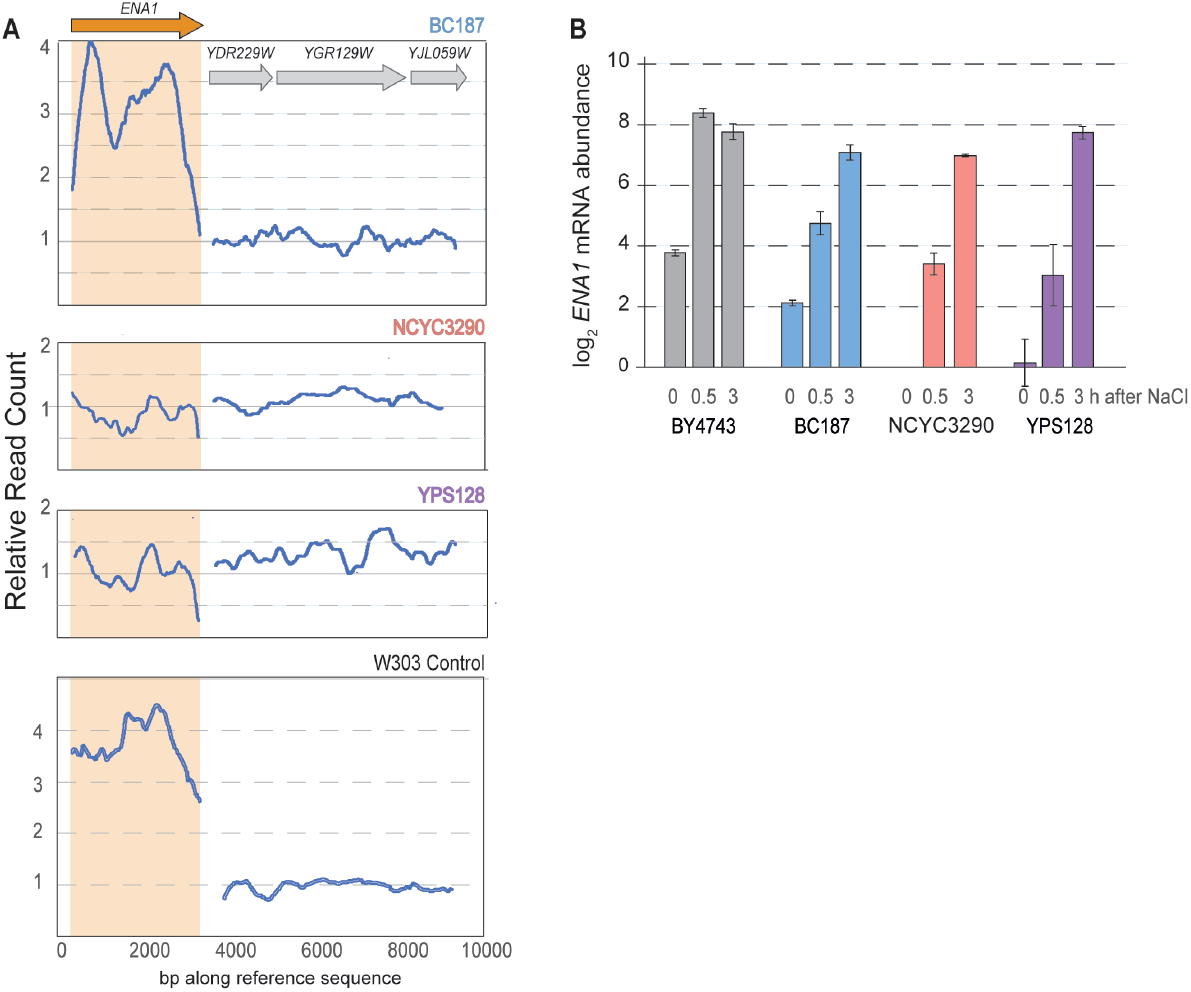
Strains vary in *ENA* copy number and expression. A) Relative DNA-seq read count across a representative *ENA* gene (left trace, orange arrow) and three representative single-gene copies (right trace, grey arrows) in each of the three wild strains (see Methods). The plot shows the running average across 500bp windows from left to right along each sequence. As a control, sequence is shown for a related laboratory strain W303, sequenced with the same pipeline as the wild strains and known to carry four *ENA* copies. B) Average and standard deviation (n=3) of log_2_ fold-change in *ENA1* mRNA abundance before and after NaCl treatment, normalized to NCYC3290 basal levels at time 0 min.

Strains with higher abundance of *ENA1* and related sodium pumps are well known to have increased tolerance to sodium stress (56, 79, 80, 84-86). Interestingly, however, strain-specific differences in *ENA* transcript abundance were not fully explained by differences in ENA copy number: although BY4743 and BC187 both harbor 3 *ENA* copies per haploid genome, basal expression in the lab strain was 3-fold higher than BC187 (Fig 4C); likewise, basal expression in BC187 was >4-fold higher than the sensitive strains. These results suggested that the response to NaCl could be influenced both by variation in *ENA* gene copy number and by *ENA* gene regulation. ENA genes are also transcriptionally induced by NaCl (84, 87, 88), and in fact several studies have observed natural variation in *ENA* transcriptional regulation (81, 89). We plotted *ENA1* mRNA abundance before and after NaCl in each strain, normalized to NCYC3290 basal levels. While BY4743 harbored the highest *ENA1* expression in the absence of stress, the strain further induced *ENA1* another 25-fold within 30 minutes after 0.7M NaCl exposure. Remarkably, the other strains all induced *ENA1* expression to within 2-fold of BY4743 mRNA levels, albeit with delayed kinetics. While induction of *ENA1* may help with the acclimation to continued salt stress (89), it is likely that the basal expression levels influence the immediate survival after rapid-onset salt stress. Thus, we propose that although strains retain the ability to induce *ENA1* expression after stress, the low starting mRNA levels likely contribute to variations in the ability to survive the initial NaCl exposure – this may also explain differences in the fitness cost of other OE genes, including OE of other sodium transporters and ion-response regulators whose duplication provides a major benefit to wild strains but has no effect in BY4743 (see Discussion).

## DISCUSSION

Our results show that environmental stress has a major influence on the fitness consequences of gene OE, and those effects vary substantially across genetically distinct individuals. That environmental stress influences the cost of gene OE may not seem surprising from a physiological point of view, and this is consistent with longstanding models on the effects of stress on mutational tolerance (90-93). However, our results have major implications for how strain and environment affect the fitness costs of gene CNV, which is an important route to rapid evolution. These trends are almost certainly true for other stresses beyond NaCl treatment studied here. We previously showed only small overlap in OE fitness consequences to toxin tolerance in wild strains growing in industrial conditions (94). We subsequently showed that the mechanisms of tolerating those conditions vary substantially across strains, suggesting that different genes will be beneficial depending on the physiological weaknesses of each individual (53). We propose that Strain-by-Environment-by-CNV interactions are prominent and could produce substantial variation in the evolutionary trajectories accessible to different individuals and over space and time.

Simply duplicating a gene’s copy number can increase its expression, at least for genes that are not dosage regulated (17, 52, 95, 96). Amplification of many genes is deleterious even in the absence of stress, likely due to the increased burden of producing extra DNA, RNA, and protein but also due to internal imbalances caused therein (16, 97). NaCl treatment exacerbated the deleterious effects of many of those genes, as might be expected (90-93). But even for the same concentration of NaCl, strains more sensitive to NaCl showed both more deleterious responses and larger effect sizes during NaCl and compared to more tolerant strains (Fig 2). This is consistent with the idea that the cost of CNV is generally worse when cells are already taxed, in accordance with previous implications from our work (18). In many cases, the increased cost of gene OE may be independent of the gene’s function (15, 16, 18) and could simply represent compounded burdens of producing extra protein during an energy-consuming stress response. In other cases, specific gene functions may be counterproductive during the NaCl acclimation and thus uniquely deleterious in that environment. For example, increased expression of functional aquaporin water transporters is detrimental during osmotic shock, due to passive water loss that exacerbates stress-induced water efflux (6). Although increased aquaporin expression is beneficial in other environments, the cost of that increase is harder to overcome in high-osmolar conditions. The implication is that different evolutionary routes will be more or less accessible depending on the environmental context.

Adaptive benefits of CNVs are well known during environmental stress (8-12), and thus we expected to find some genes whose OE is uniquely beneficial during NaCl exposure compared to standard conditions. The surprise was that there was little overlap in beneficial genes across strains, including the more sensitive strains. Part of the low overlap results from inclusion of the laboratory strain, BY4743. This strain showed the fewest beneficial genes, and most of those genes produced only mild benefits. This along with growth responses shown in Fig 1 are consistent with the notion that BY4743 is fairly tolerant of NaCl, and thus there is little room for improvement. Our genomic analyses raised several possible explanations. Unstressed BY4743 shows higher expression of many genes directly related to stress defense (Fig 4B,C), including genes linked to osmotic and other stress defenses. Furthermore, BY4743 displays very high expression of ENA genes encoding sodium efflux pumps even in the absence of stress, likely due to both natural ENA gene amplification and higher expression from one or more copies. Together, these results strongly suggest that BY4743 is already prepared for NaCl stress before exposure, requiring less acclimation effort upon treatment. This preparedness could also explain the accelerated transcriptomic response to NaCl shock (Fig 4A,B): higher basal stress tolerance coupled with immediate Na+ efflux could result in a lower effective dose of NaCl, which is known to produce a faster signaling response (64-67). Whether these expression differences have been selected due to laboratory domestication or merely accumulated due to neutral drift, these results are consistent with the notion that there is little adaptive benefit for most OE genes in the lab strain under these conditions.

The results in the lab strain are perhaps not surprising, since laboratory strains are often outliers in their responses (52, 98-100) – but we were surprised to see the low overlap in beneficial OE genes across wild strains, especially NaCl-sensitive NCYC3290 and YPS128 (Fig 3B). The beneficial gene sets for each strain were enriched for distinct functions; the exception was shared enrichment of osmo-responsive Sko1 targets among the beneficial genes shared by the two sensitive strains. Yet there were several genes common to two or all three of the wild strains (albeit with different effect sizes, often greatest in the sensitive strains, Dataset 2). These included genes linked to cation stress, including kinases Hal5 and Sat4 and transporter Qdr1 that are known to modulate ion tolerance (60, 101, 102). It is interesting that these genes had virtually no benefit in the lab strain that is already well equipped for Na+ stress.

We propose that the significant fitness benefit of these genes to wild strains but not BY4743 could be impacted by variation in abundance of *ENA* ATPase Na+ pumps. The sensitive strains harbor only one copy of *ENA* per haploid genome, whereas BC187 carries 3 copies that remain lower expressed than in BY4743 (Fig 5). Given the importance of ENA expression in NaCl tolerance (79, 80), we propose that differences in *ENA* copy number and expression influence which other OE genes will be beneficial. This would explain why strains with lower *ENA* mRNA abundance greatly benefit from OE of other genes directly involved in sodium efflux, whereas BY4743 receives no benefit from over-expression of those genes. Interestingly, several other genes uniquely beneficial to one or more wild strains transcriptionally up-regulate *ENA1*, including CK2 whose OE benefited all strains and *CRZ1* that was highly beneficial to BC187 (37, 84, 103). These possibilities highlight the potential for genetic interactions between CNVs, since the impact of one gene’s amplification is dependent on another gene’s copy number.

In all, this study adds to the body of evidence that the impact of a mutation, in this case gene OE to mimic the effects of CNV, depends on not only the genomic context but also environment. Ultimately, the combined effect of genomic and environmental variation can perhaps best be considered as Gene-by-System interactions (104), where the impact of a gene’s CNV depends on the physiological state of the cell. A remaining challenge is developing statistical models that can both represent Gene-by-System interactions and uncover the biology behind them.

## ACKNOWLEDGEMENTS

We thank members of the Gasch Lab for helpful discussions. This work was funded by R01 GM147271 to APG and in part by a grant to the Great Lakes Bioenergy Research Center from the UW Department of Energy (DE-S0018409).

## DATASET LEGENDS

**Dataset 1: Normalized and imputed barcode read count**. Normalized and imputed data from MoBY 2.0 selection experiments, as described in Methods.

**Dataset 2: *EdgeR* output for MoBY 2.0 selections**. Each block represents edge R results for each indicated strain, see *edgeR* manual for details.

**Dataset 3: Beneficial genes**. Each tab represents *edgeR* output for genes determined to be significant at FDR < 0.05. Tabs for BY4743 and BC187 labeled ‘threshold’ are those with FDR < 0.05 and meeting the effect size threshold as described in Methods.

**Dataset 4: Normalized log2 relative gene expression from RNA-seq experiments**. Tab 1) log_2_ values represent the fold change in expression in each denoted strain at 30 min or 3 h versus the unstressed sample for that strain. Tab 2) log_2_ values represent the fold difference in expression in each denoted unstress strain relative to the mean expression of all four strains from that paired replicate.

